# The evolution of realized niches within freshwater *Synechococcus*

**DOI:** 10.1101/678375

**Authors:** Nicolas Tromas, Zofia E. Taranu, Mathieu Castelli, Juliana S. M. Pimentel, Daniel A. Pereira, Romane Marcoz, B. Jesse Shapiro, Alessandra Giani

## Abstract

Understanding how ecological traits have changed over evolutionary time is a fundamental question in biology. Specifically, the extent to which more closely-related organisms share similar ecological preferences due to phylogenetic conservation – or if they are forced apart by competition – is still debated. Here we explored the co-occurrence patterns of freshwater cyanobacteria at the sub-genus level to investigate whether more closely-related taxa share more similar niches, and to what extent these niches were defined by abiotic or biotic variables. We used deep 16S rRNA gene amplicon sequencing and measured several abiotic environmental parameters (nutrients, temperature, etc.) in water samples collected over time and space in Furnas Reservoir, Brazil. We found that relatively more closely-related *Synechococcus* (in the continuous range of 93-100% nucleotide identity in 16S) had an increased tendency to co-occur with one another (*i.e.* had similar realized niches). This tendency could not be easily explained by shared preferences for measured abiotic niche dimensions. Thus, commonly measured abiotic parameters might not be sufficient to characterize, nor to predict community assembly or dynamics. Rather, co-occurrence between *Synechococcus* and the surrounding community (whether or not they represent true biological interactions) may be a more sensitive measure of realized niches. Overall, our results suggest that realized niches are phylogenetically conserved, at least at the sub-genus level and at the resolution of the 16S marker. Determining how these results generalize to other genera and at finer genetic resolution merits further investigation.

**Originality-Significance Statement:** We address a fundamental question in ecology and evolution: how do niche preferences change over evolutionary time? Using time-series analysis of 16S rRNA gene amplicon sequencing data, we develop an approach to highlight the importance of biotic factors in defining realized niches, and show how niche preferences change proportionally with the 16S gene molecular clock within the genus *Synechococcus.* Ours is also one of few studies on the ecology of freshwater *Synechococcus*, adding significantly to our knowledge about this abundant and widespread lineage of Cyanobacteria.

## Introduction

A bacterial community is a group of potentially interacting organisms that coexist at a particular place and time (Magurran, 2003). Environmental selective pressures are a strong force shaping microbial community assembly (Martiny et al, 2015). We know, for example, that certain abiotic factors explain a large portion of the variation in microbial community composition (e.g. the effect of pH on soil bacterial communities; Fierer and Jackson, 2006). Therefore, associations between microbial traits - generally defined as a phenotypic response to a specific environmental condition - and abiotic niches could help explain community assembly rules (Green et al., 2008; Burke et al., 2011). However, using trait-based approaches to understand communities may be challenging, as important abiotic variables may go unmeasured. Even less is known about biotic interactions, despite their importance in determining community composition, diversity, and dynamics (Needham et al., 2016).

Abiotic factors are generally thought to determine an organism’s fundamental niche (where it is theoretically capable of living), whereas biotic factors determine its realized niche (where it actually lives in nature; Hutchinson, 1957). Species (or taxonomic) co-occurrence networks are often used to infer niche similarity among organisms that tend to co-occur in nature over space and time, and microbial co-occurrence networks can easily be constructed from 16S rRNA gene amplicon sequencing surveys (Friedman and Alm, 2012; Röttjers and Faust, 2018; Tromas et al., 2018). However, co-occurrence can be driven by both abiotic and biotic factors, which are hard to disentangle in practice (Kraft et al., 2015). Regardless, organisms sharing a similar niche are expected to be associated with similar surrounding communities (Cohan and Koeppel, 2008; Faust et al.2012; Pascual-García et al., 2014).

A fundamental question spanning ecology and evolution is how ecological traits change over evolutionary time. For example, some traits (such as bacteriophage host range or phosphorus utilization) evolve rapidly at the tips of a phylogenetic tree, whereas other traits (such as salinity preference) are deeply conserved (Martiny *et al.* 2015). When more closely related organisms share similar ecological preferences, so-called “habitat filtering” or “environmental” filtering is expected to result in phylogenetic clustering, meaning that a community tends to contain more closely related organisms than expected by a random draw from the phylogeny (Webb et al., 2002; Horner-Devine & Bohannan, 2006; Martiny et al., 2015). In contrast, if close relatives evolve different traits to avoid competitive exclusion, this will result in phylogenetic overdispersion (*i.e.* a community composed of more distant relatives than expected by chance). However, the relative importance of these two processes in shaping microbial communities is still widely debated, and difficult to distinguish (Koeppel and Wu, 2014; Cadotte and Tucker, 2017). We have previously shown that within the cyanobacterial genus *Dolichospermum* (Tromas et al., 2018), the relationships between phylogenetic distance and ecological similarity varies by trait, suggesting that it might be necessary to analyze each niche dimension or trait separately (Martiny *et al*., 2015).

In this study, we explored the co-occurrence patterns of freshwater cyanobacteria at the sub-genus level, to investigate if more closely related taxa have more similar niches, and to what extent these niches can be quantified by abiotic or biotic variables. We use the term “closely related” as a relative term, describing a continuous range of genetic similarity (93-100% nucleotide identity) in 16S rRNA gene sequences. We focused on *Synechococcus*, the most abundant cyanobacterial genus in Furnas Reservoir (Brazil) at the time of sampling (2006-2008). *Synechococcus* is among the most abundant organisms living in oceans and lakes (Stockner et al. 2000; Scanlan, 2003). The phylogenetic coherence of the genus has been questioned by a recent study showing it to be polyphyletic (Coutinho et al 2016). *Synechococcus* is physiologically highly plastic, ubiquitous, and able to acclimate to different environmental conditions (Callieri 1996; Vörös et al. 1998; Callieri et al., 2011). Previous studies have shown that different *Synechococcus* strains could co-exist in the same site but respond differently to environmental changes, suggesting niche partitioning (Ferris et al., 2003; Allewalt et al., 2006; Becker et al., 2007; Becraft et al., 2011; Callieri et al. 2012). Recently, Zheng et al. (2018), observed a geographical pattern of heterotrophic bacteria associated with different marine *Synechococcus* strains, indicating that strains living in the same area tend to be associated with similar communities. It remains, however, unclear whether the *Synechococcus* genotype or the environment are the main drivers of *Synechococcus* interactions with the surrounding microbial community.

Using deep 16S rRNA gene amplicon sequencing of 86 water samples collected in time series across nine locations in the Furnas Reservoir, we tracked genetic diversity within the *Synechococcus* genus, along with the surrounding microbial community, and measured several abiotic variables. We found that that more closely-related *Synechococcus* tended to co-occur with one another and also with similar surrounding microbial communities. Such phylogenetic clustering indicates that realized niche similarity tends to evolve proportionally with substitutions in the 16S gene in the *Synechococcus* lineage (*i.e.* niches evolve according to the 16S gene molecular clock). However, more closely-related *Synechococcus* did not have similar abiotic niche preferences (with the exception of total phosphorus). These results suggest that biotic factors may be strong niche determinants, which could be particularly useful when relevant abiotic factors are unknown or cannot be readily measured. Alternatively, cryptic abiotic drivers may determine niche and community structure, but biotic factors provide the most informative measure of niche similarity.

## Materials and methods

### Sampling, environmental data measurements

A total of 90 water samples were collected from September 2006 to April 2008 at nine stations in Furnas Reservoir (Minas Gerais, Brazil). Furnas is a large reservoir (1440 km^2^), located in southeastern Brazil (20°40’S; 46°19’W) and formed by the damming of two main rivers (Rio Grande and Rio Sapucaí), which divide the reservoir in two separated branches (Figure S1). Sampling stations F1, F4, F6, F9 are from a relatively pristine branch of the reservoir, whereas F12, F14, F18 and F20 are impacted by human activities. Temperature profiles were measured in the water column by aid of a multi-parameter probe (556 YSI, USA). Water samples were collected from the euphotic zone (determined by Secchi disc depth) by a Van Dorn vertical water sampler. Samples were stored in bottles that had been acid-washed and rinsed with deionized water. A portion of each water sample was immediately filtered through glass fiber filters (Whatmann GF/F, 0.7 μm pore size). The exact filtered volumes were recorded and filters were kept frozen until further analyses (chlorophyll a, DNA and microcystin). For dissolved nutrient analyses, 200 mL of filtered water samples were stored at −20 °C. For total phosphorus, samples were frozen with no previous filtration. Nutrient analyses were performed using spectrophotometric methods according to APHA (2005). All nutrient analyses (Nitrate, Nitrite and total phosphorus (TP) were performed on three replicates.

### DNA extraction, purification and sequencing

DNA was extracted from frozen filters according to Kurmayer et al. (2003), with few modifications. Briefly, filters were treated with a sucrose buffer (25% w/v sucrose, 50 mM Tris-HCl, 100 Mm EDTA, pH 8) on ice for 2 h and with addition of lysozyme (5mg/mL, 1h, 37°C). Proteinase K (100 μg/mL) in sodium dodecyl sulfate (2% v/v) was added and filters were incubated overnight at 55°C. A phenol:chloroform:isoamyl alcohol solution (25:24:1, v/v) was used for protein precipitation and DNA isolation. The DNA was cleaned in 100% ethanol and pellets rinsed with 70% ethanol. The DNA was resuspended in TE (10 mM Tris-HCl, pH 8, and 1 mM EDTA). The DNA extract was quantified by a spectrophotometer (Perkin Elmer, Lambda 25), at 260 nm and 280 nm, and its quality checked in 1% (w/v) agarose gel, stained with ethidium bromide. DNA samples were stored at −20°C.

### Sequence analysis

We followed the same protocol described in Tromas et al., (2017) to generate a library of V4 region amplicons. Libraries for 2×250bp paired-end Illumina sequencing were prepared using a two-step 16S rRNA gene amplicon PCR as described in Preheim et al. (2013). We performed one sequencing run using MiSeq reagent Kit V2 (Illumina, San Diego, CA, USA) on a MiSeq instrument (Illumina). A total of 4,476,747 sequences of the 16S rRNA gene V4 region were obtained from 90 lake samples, two negative controls, and two mock community samples. We obtained a median of 37,682 sequences per sample. Using a similar approach as described in Tromas *et al*., (2018), we processed the sequences with SmileTrain (https://github.com/almlab/SmileTrain/wiki; Preheim et al., 2013) for read quality filtering, primer removal, chimera filtering, and merging using USEARCH (version 7.0.1090, http://www.drive5.com/usearch/, default parameter) (Edgar, 2010), Mothur (version 1.33.3) (Schloss *et al*., 2009), and Biopython (version 2.7). Minimum Entropy Decomposition (MED) was then applied to the filtered and merged reads to partition sequence reads into MED nodes (Eren *et al*., 2015). MED was performed using the following parameters: –M noise filter set to 1000, resulting in ∼17.5% of reads filtered and 466 MED nodes representing the whole bacterial community. Samples with less than 1000 reads were removed, yielding a final dataset of 86 reservoir samples. Finally, we assigned taxonomy to MED nodes using the assign_taxonomy.py QIIME script (default parameters), and a combination of GreenGenes and a freshwater-specific database (Freshwater database 2016 August 18 release; Newton *et al*., 2011), using TaxAss (https://github.com/McMahonLab/TaxAss, installation date: September 13^th^ 2016; Rohwer *et al*., 2017). We found a total of 40 *Synechococcus* nodes, which we refer to as strains below.

### Spatio-temporal analysis

We used multivariate regression tree analyses (Breiman *et al*. 1984; De’ath 2002) to investigate if spatio-temporal variables could explain genetic variation within *Synechococcus*. We used two different temporal predictors: year and month and one spatial predictor: station. The analysis was performed using the function *mvpart*() and *rpart.pca*() from the R mvpart package (Therneau and Atkinson, 1997; De’ath, 2007). Prior to analysis, the *Synechococcus* MED node table was Hellinger transformed to downweight the effect of double-zeros (Rao, 1995).

### Genetic distance between Synechococcus nodes

We measured the genetic distance between *Synechococcus* MED nodes using the software MEGA (Kumar et al., 2016; version 7.0.18) with the p-distance (the proportion of nucleotide sites at which two sequences differ), calculated by dividing the number of sites with nucleotide differences by the total number of sites compared (excluding sites with gaps).

### Co-response to abiotic factors

As described in Tromas *et al.* (2018), we used a Latent Variable Model (LVM) framework (boral package in R; Hui 2015, Warton *et al.* 2015) to explore how *Synechococcus* nodes co-responded to abiotic gradients and used these co-responses as indicators of niche similarity. That is, for each abiotic factor, we ran separate LVMs, regressing the bacterial community as a function (both linear and non-linear) of the given factor. This component of the LVM thus defined the taxon-specific environmental responses. To then identify remaining patterns of co-occurrence after accounting for all measured environmental variables, we fit a global LVM which included all abiotic factors and two latent variables (*sensu* Letten *et al.* 2015 and Warton *et al.* 2015). To visualize patterns of co-occurrence arising from the different environmental factors, we calculated two types of correlation matrices. The first, a co-response correlation matrix, was constructed by calculating, for any two nodes, the correlation between their fitted values. This correlation matrix thus represented the correlation between nodes that can be attributed to a shared or diverging environmental responses. A significant positive correlation between the fitted response of any two taxa to an environmental variable represented a co-response (i.e., as one taxa increases in response to an environmental variable, the other likewise increases), whereas a significant negative correlations between the fitted response of two taxa represented some degree of niche separation (i.e., as one taxa increases in response to an environmental variable, the other decreases).

To account for the correlation between nodes that may be attributable to biotic processes or missing environmental covariates, a second type of correlation matrix, a residual correlation matrix, was calculated using the latent variable coefficients of the global LVM. Since the latent variables are the output of an ordination of the residuals, we expected weak residual correlations induced by the latent variables if the environmental variables were the dominant force structuring patterns of species co-occurrence (*i.e.* the model explained most of the variance, with little left unexplained in the residuals). Conversely, if any unmeasured environmental factors and/or biotic processes are equally, or more, important than measured environmental factors, we expected strong correlations based on the latent variables (*i.e.* the model was poor and much of the observed variability remained in the residuals).

To test how *Synechococcus* niche similarity varied with genetic distance, we examined the relationship between the co-response of *Synechococcus* taxa to environmental parameters and their genetic distance (i.e., plotting the correlation coefficient of the LVM co-responses vs. genetic distances) (R_script1). However, given the large number of environmental factors driving phytoplankton community dynamics (Hutchinson, 1961), we expected some degree of co-limitation whereby the response of a taxon to an environmental variable (and consequently the degree of co-response among any two taxa) would be limited by other environmental variables. In such cases, the taxon’s response and rate of change would have an upper limit (set by all measured environmental factors) but may not reach this limit if other, unmeasured factors are co-limiting (Cade and Noon 2003). As an increasing number of unmeasured factors become limiting at some sample location, or time point, the relationship between the response and the measured factor becomes increasingly heterogeneous or wedge shaped. When such heterogeneous variances are observed (wedge shape biplot), it suggests that there is not a single slope coefficient that characterizes the relationship, and that focusing solely the coefficient fit by ordinary least squares regression (mean response) may underestimate the true rate of change. Thus, to accurately examine co-limitations among measured and unmeasured factors, we applied quantile regressions using ‘quantreg’ R package (Koenker, 2015) (R_script1), which, instead of fitting a regression to the mean response (ordinary least squares regression), fits regression curves to other quantiles of the response variable (Cade and Noon 2003). Finally, to provide an effect size for the linear quantile relationships of abiotic co-responses vs. genetic distance, we calculated the goodness of fit measure for quantile regression for a suite of quantiles (where the goodness of fit is estimated as 1 minus the ratio between the sum of absolute deviations of the parameterized model and the sum of absolute deviations of the unparameterized, null model). Note that this goodness of fit measure (referred hereafter as R1) is not comparable to the OLS coefficient of determination (R2); it is based on the variance of absolute deviations as opposed to the variance of squared deviations, and consequently, R1 is always smaller than R2 (Koenker and Machado, 1999).

### Co-occurrence with biotic factors

A caveat of the above LVM-quantile regressions is that although it identifies whether some relationships are limited by unmeasured abiotic and/or biotic factors, it could not tease apart the relative importance of each type of factor (everything is instead lumped as an unknown). In order to determine the relative contribution of biotic processes, we thus quantified node co-occurrence patterns among *Synechococcus* and the remaining community. In particular, we explored the relationship between *Synechococcus* nodes and the surrounding community by measuring: (1) co-occurrences between *Synechococcus* and other taxa, and (2) paired differences in co-occurrence among *Synechococcus* nodes and a specific taxon. For the former, we calculated co-occurrences among taxa using SparCC (Friedman and Alm, 2012), including 20 iterations to estimate the median correlation of each pair of MED nodes, 500 bootstraps to assess the statistical significance and centered log ratio (CLR) transform to correct for compositionality. Correlations were then filtered using a false discovery rate threshold (*Q* < 0.05). For the latter, we further calculated the absolute difference of SparCC correlation (*r*) between a *Synechococcus* node (X_i_) and a specific taxon T, such that |Δ*r*| = | Corr(X_1_, T) - Corr(X_2_, T) | where Corr is defined here as the SparCC correlation between X_i_ and T.

For each non-*Synechococcus* taxon, we then estimated the relationship between |Δ*r*| and the distance between the given *Synechococcus* nodes (X_1_ and X_2_). A positive correlation would indicate that more closely related *Synechococcus* nodes have a lower |Δ*r*| than more distant nodes, *i.e.* more closely related nodes would have more similar correlations with potentially interacting community members. To reduce potential bias or noise due to a small sample size of *Synechococcus* pairs, we selected all non-*Synechococcus* nodes that were significantly co-occurrent with at least seven different *Synechococcus*. This cutoff was chosen to ensure at least 20 unique pairs of *Synechococcus* for each non-*Synechococcus* taxon. A total of 373 non-*Synechococcus* taxa were selected using this cutoff, and the correlation between |Δ*r*| and genetic distance was non-significant (Spearman correlation, P<0.05) for 120 of them. Finally, we performed a permutation test to quantify any bias in the method due to data structure. For each of the significant non-*Synechococcus* taxa, we estimated the rate of false positive correlations by sampling values of *Synechococcus* genetic distance with replacement, while randomizing the association between genetic distance and |Δ*r*|. We performed 1000 permutations for each significant non-*Synechococcus* taxon using the script R_script2. We then recorded the proportion (*p*) of permutations yielding a larger correlation than observed, after adding a pseudocount of 1 to both the numerator (the number of permutations yielding a larger correlation than observed) and the denominator (the total number of permutations).

### Phylogenetic analysis

We first verified that our *Synechococcus* sequences were monophyletic by building a phylogenetic tree with FastTree (version 2.1.8, Price et al., 2009), using all MED node sequences aligned with MAFFT (v7.154b; Katoh and Standley, 2013). We then specifically aligned *Synechococcus* MED node sequences using muscle (in Mega, version 7.0.26; Kumar et al., 2016) and built phylogenetic trees using PhyML with the general time-reversible (GTR) model of nucleotide substitution (version 3.0; Guindon et al., 2010). We used the ALDEx2 R package (version: 1.5.0; Fernandes *et al*., 2014) and the aldex() function to identify MED nodes associated with the pristine or human-impacted branches of Furnas Reservoir. To do so, we used Welsh’s t-test and 128 Monte Carlo samples. ALDEx2 uses the CLR transformation to avoid compositional effects. A Q-value below 0.05 after Benjamini-Hochberg correction was taken as evidence for association. We also used a second association method, DESeq2 implemented in the “MicrobiomeAnalyst” web-based tool (Dhariwal et al., 2017).

### Statistical analysis

All statistical tests were performed in R.

## Results

### Abiotic factors shape Synechococcus community structure

Furnas Reservoir is divided into two branches: one, which is heavily human-impacted (Figure S1, sampling stations 12-20) and the other, less impacted (stations 1-9). *Synechococcus* dominated the bacterial community in both branches during the sampling period (2006-2008) (Figure S2) and formed a monophyletic group (Figure S3, S4). Furnas Reservoir is meso-eutrophic based on phosphorus concentrations. Total phosphorus (TP) did not vary significantly across stations but did vary significantly within each station over time (Figure S5). Temperature did not vary widely across sampling stations or time. We first asked whether particular *Synechococcus* strains (which we used interchangeably with MED nodes) were associated with different branches of the reservoir, different time periods, or different abiotic factors. We applied a multivariate regression tree analysis (Breiman et al. 1984; De’ath 2002) to test whether the composition of *Synechococcus* strains changed over time and space. We found that the year and month of sampling did not explain much variance (R^2^=0.02 and R^2^=0.03, respectively), nor did the station (R^2^=0.06) or branch of the reservoir (R^2^=0.02). In contrast, Nitrite, TP, depth, and water temperature all explained a significant proportion of the variance (Figure S6; 22.5% of variance explained). Nitrite is the single most explanatory variable (13% of variance, but also colinear with temperature and depth; Figure S6), followed by TP (5.5% of variance; Figure S7).

### More closely related Synechoccus do not share similar preferences for most measured abiotic niches

Having established that abiotic factors, but not temporal or spatial variation, explain a significant amount of variation in the relative abundances of *Synechococcus* strains, we asked whether more closely related strains tended to share similar abiotic niches. We examined *Synechococcus* 16S sequences across a range of ∼93-100% nucleotide identity, which is a rather large range considering that niche partitioning can occur at relatively high identity, >97% (Martiny *et al*. 2015; Larkin and Martiny, 2017). Here we defined abiotic niche similarity by estimating the co-responses of pairs of *Synechococcus* strains to each abiotic factor using LVM-quantile regression (Methods). For most abiotic factors, we did not observe any significant relationship between co-responses and genetic distance, with the exception of total phosphorus for which the relationship was negative (Median p-value across quantiles = 0.021; median R1 across quantiles = 0.12) (Figure 1 & S8, Table S2). Similarly, the degree of abiotic niche separation (*i.e.* significant negative correlations between the fitted response of any two *Synechococcus* strains to an abiotic factor) was generally independent of genetic distance (Figure S9, S10), except for depth and Nitrite which were significant at some quantiles but with quite a poor fit (median R1 across quantiles ≤ 0.02)(Table S2C).

**Figure 1.**
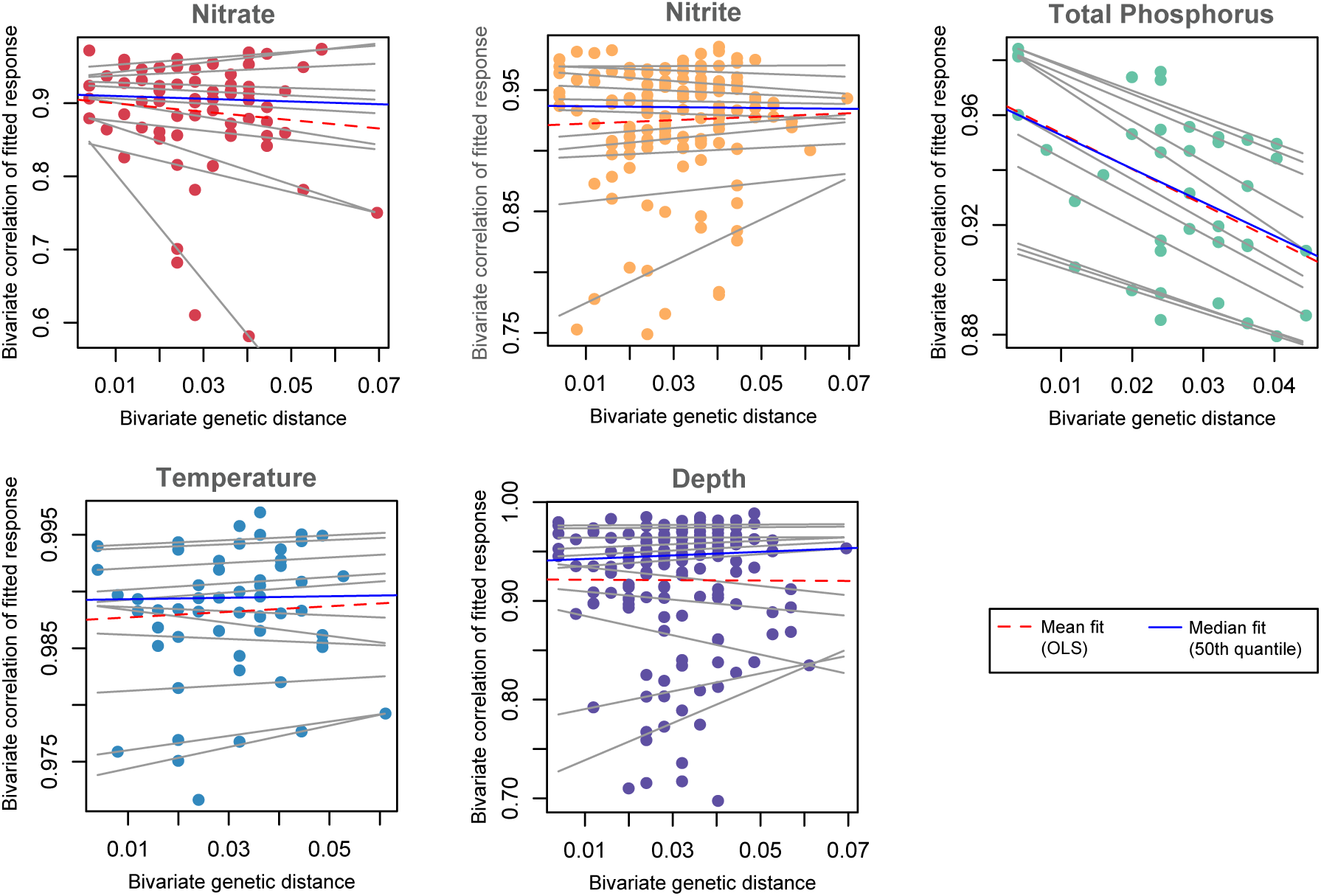
Relationship between *Synechococcus* niche similarity and genetic distance. Niche similarity is defined as a positive LVM co-response (y-axis). Shown are the scatterplots of LVM co-responses versus each variables superimposed with the lines for 5^th^, 10^th^, 15^th^, 25^th^, 35^th^, 45^th^, 55^th^, 65^th^, 75^th^, 85^th^ and 90^th^ quantile regression fits (grey lines, for each co-response quantile), the median fit (50^th^ quantile; blue line), and the least squares estimate or mean fit (dashed red line).

We also evaluated non-linear quantile relationships among co-responses and genetic distances, but we never found a significant non-linear relationship over the total range of genetic distances (Table S2A-C). We repeated the quantile regression analysis in a restricted range of 100-97% identity, to see if any linear trend is discernible in a range roughly corresponding with species or genera (Figure S11). The negative linear trend observed between co-responses and genetic distance for TP in the full range of genetic identify (Figure 1) was still observed in the restricted range (Figure S11) although not significant at all quantiles (Table S2D). There was also some evidence for a non-linear relationship between genetic distance and both positive and negative co-responses to TP (Table S2 D, E), suggesting possible niche separation among closely related (∼99% identical) strains. However, this trend is driven by just three pairs of very closely related strains with quite different TP preferences (Figure S12). Overall, these results suggest that TP-related niches show some evidence for phylogenetic conservation within freshwater *Synechoccus*, while the evidence for other niches is weak or inconclusive.

Biotic processes or unmeasured environmental variables could define *Synechococcus* realized niches, and are captured in the residual variation of the global LVM. We next asked whether more closely-related *Synechococcus* strains had more similar co-responses to this residual variation. For the positive co-responses (Figure 2A, B), we observed a decrease with genetic distance. This result suggests a negative relationship between the residuals (*i.e.* unmeasured abiotic variables and/or biotic niche preferences) and phylogenetic distance. For the negative residual correlations (Figure 2C, D), there was no apparent relationship with genetic distance. Therefore, there remain significant co-responses that were not explained by the abiotic factors measured in this study. Overall, most of the measured abiotic environmental variables were not related to *Synechococcus* genetic distance. However, preferences for unmeasured biotic or abiotic factors do tend to change with genetic distance, such that more closely-related *Synechococcus* have similar realized niche preferences.

**Figure 2.**
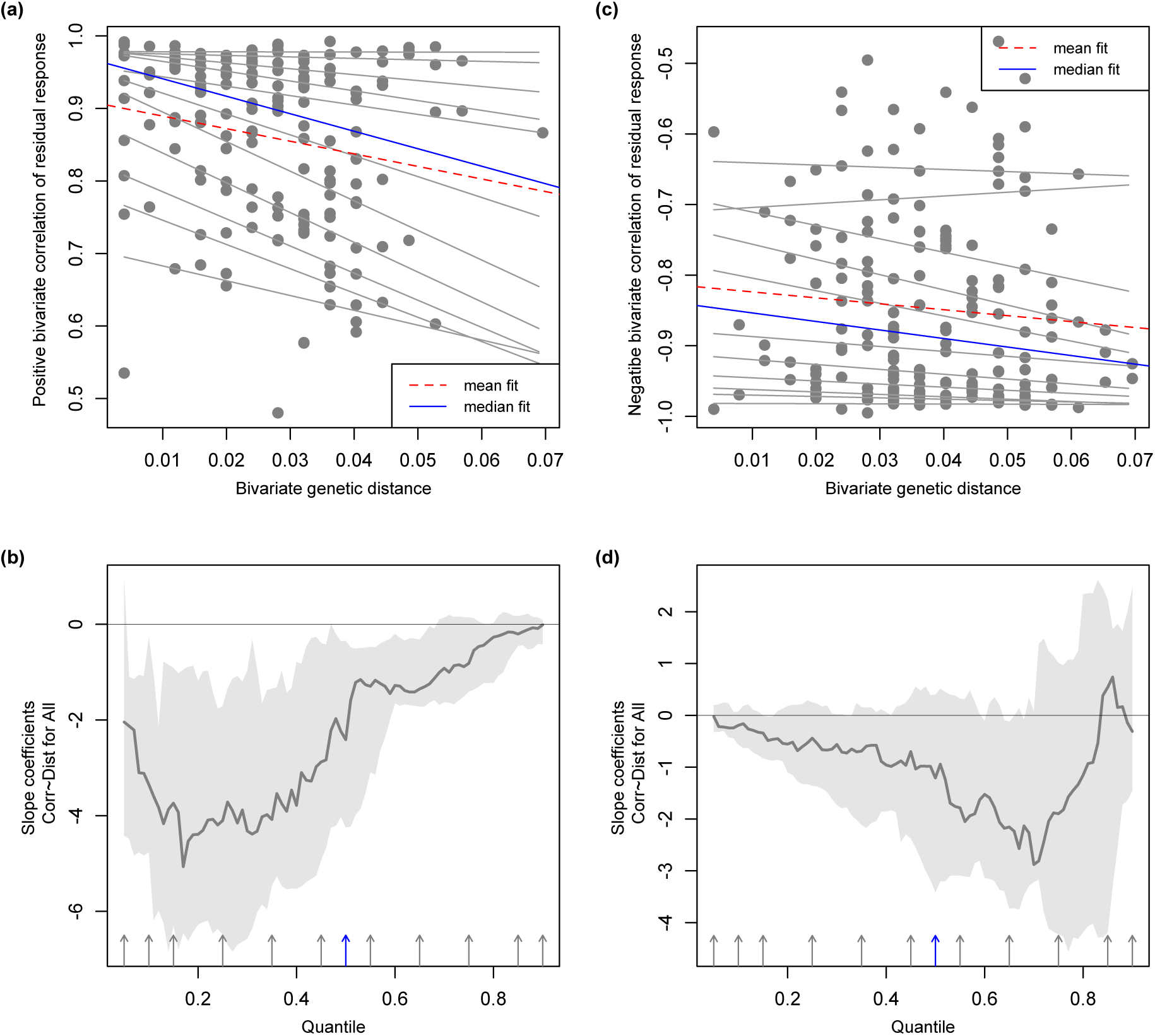
Relationship between *Synechococcus* co-responses (LVM residuals) and genetic distance. Positive co-responses are shown in panels **A** and **B**; negative co-responses in panels **C** and **D.** (A, C) Scatterplots of LVM co-responses versus LVM residuals superimposed with the lines for 5^th^, 10^th^, 15^th^, 25^th^, 35^th^, 45^th^, 55^th^, 65^th^, 75^th^, 85^th^ and 90^th^ quantile regression fits (grey lines), the median fit (50^th^ quantile; blue line), and the least squares estimate or mean fit (dashed red line). (B, D) Panels show the quantile slope estimates (dark grey line) and corresponding confidence intervals (light grey bands) of the response models across all quantiles (arrows, with the median shown in blue). Significant quantile slopes occur when confidence intervals do not overlap zero.

### More closely related Synechococcus share similar biotic interactions

The observed tendency for more closely-related *Synechococcus* to share similar realized niches (Figure 2A) could be explained by shared preferences for either unmeasured abiotic factors or biotic interactions. We used co-occurrence network analysis to explore the role of potential biotic interactions. First, we confirmed that more closely related *Synechococcus* tend to co-occur across samples. We observed a negative relationship between node co-occurrence and pairwise genetic distance (Figure 3), indicating that genetically similar *Synechococcus* nodes are indeed more likely to co-occur, and thus to have similar realized niches. This pattern was significant [linear regression, F(1,488) = 108.6, *P* < 0.001, adjusted R^2^ =0.18].

**Figure 3.**
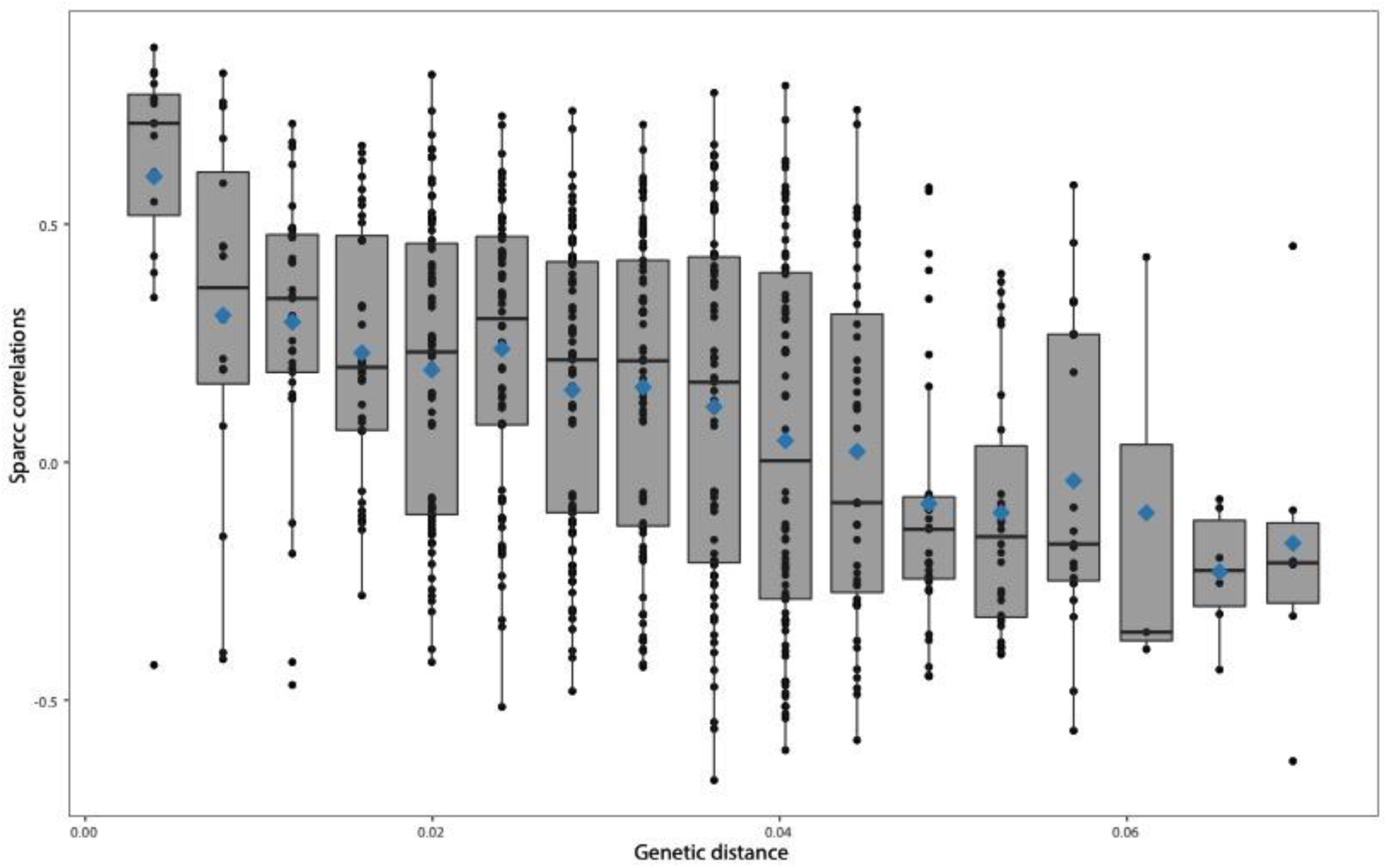
Negative relationship between pairwise SparCC correlation coefficients and pairwise genetic distance between *Synechococcus* nodes. Only significant SparCC correlations (Q < 0.05) were included. Pairwise genetic distance was computed as percent nucleotide identity (p-distance). Blue diamonds represent the mean SparCC correlation for each distance. Boxplots show the median (horizontal line), the 25th and 75th percentile (enclosed in box) and 95% confidence intervals (whiskers).

If more closely-related *Synechococcus* tend to co-occur, it follows that they should also co-occur with more similar surrounding microbial communities. The composition of the surrounding microbial community could provide information about the nature of the realized niche, while also suggesting possible biotic interactions. To confirm the expectation that more closely-related *Synechococcus* should also share similar surrounding communities, we performed a pairwise analysis to examine the relationship between *Synechococcus* genetic distance and |Δ*r*|, a measure of the similarity of *Synechococcus* co-occurrence with non-*Synechococcus* taxa (Methods). A total of 373 non-*Synechococcus* taxa were analysed and the correlation between |Δ*r*| and genetic distance was non-significant for 120 of them. The remaining 253 significant non-*Synechococcus* have consistently positive relationships between co-occurrence and genetic distance, *i.e.* more closely related *Synechococcus* nodes have a higher chance of co-occurring with similar non-*Synechococcus* taxa (Figure 4). This result is robust to data structure, as determined by permuting the associations between genetic distance and |Δ*r*| (Figure S13; Methods).

**Figure 4.**
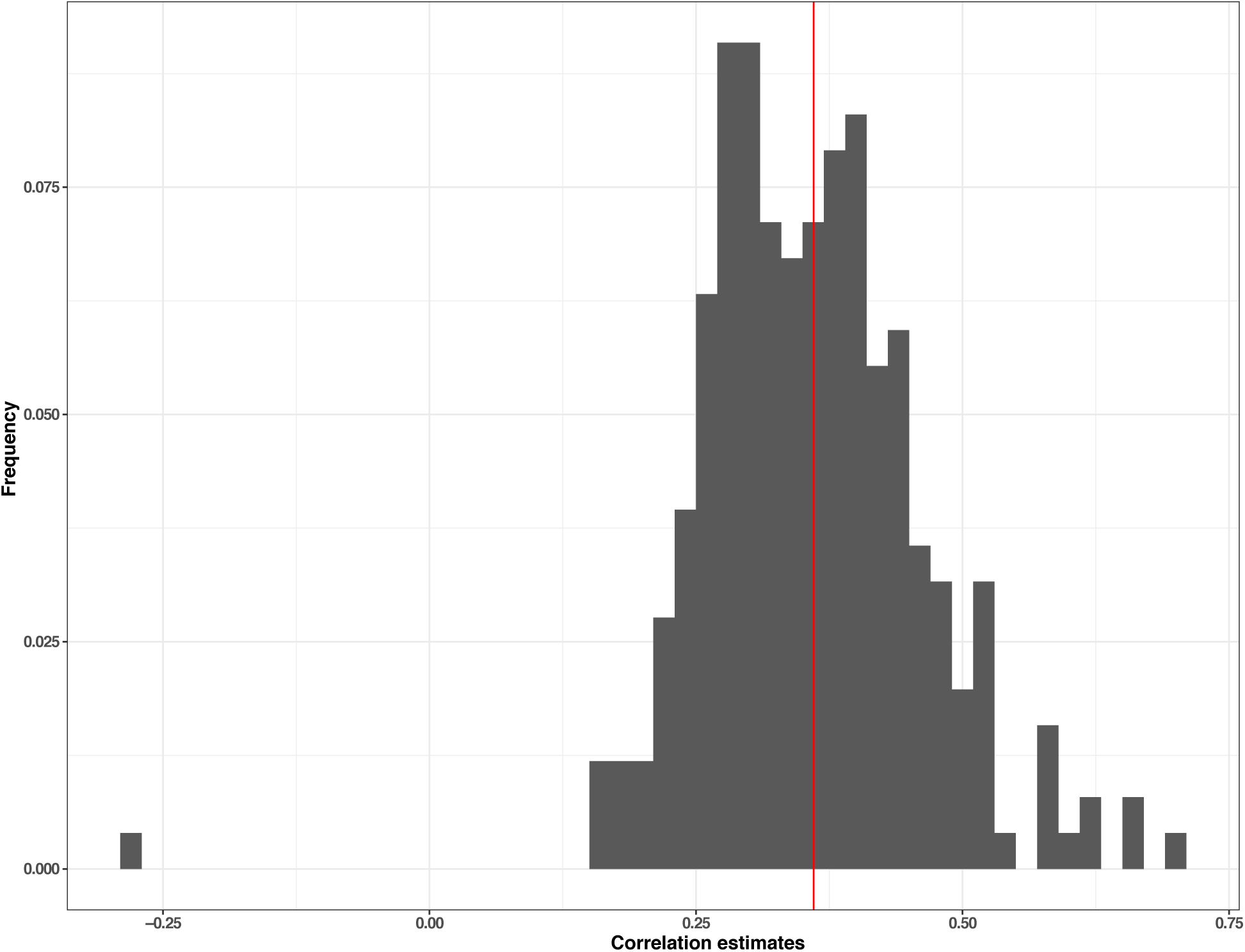
More closely related *Synechococcus* nodes co-occur with similar surrounding community. The histogram shows the distribution of Spearman correlations between the genetic distance of *Synechococcus* nodes and their association with similar surrounding communities. Spearman correlations were calculated between the genetic distance of *Synechococcus* pairs and the absolute difference of SparCC correlation (r) of a *Synechococcus* node with a specific taxon T, such that |Δr| = | Corr(X_1_, T) - Corr(X_2_, T) | where Corr is defined here as the SparCC correlation score of X_i_ (*Synechococcus* node) and T (non-*Synechococcus* node). We then estimated the correlation of |Δr| and the genetic distance between nodes X_1_ and X_2_. The mean Spearman correlation estimate was 0.373 (red line) and 99.6% of correlations are positive.

Finally, we selected the non-*Synechococcus* taxa with the highest correlation scores between |Δr| and *Synechococcus* distance genetic. Five taxa had correlation estimates higher than 0.6, including three Proteobacteria, one Verrucomicrobia, and one Planctomycetes (Table S1). Previous studies have shown that members of the *Comamonadaceae* family and *Spirobacillales* order (members of Proteobacteria; Table S1) were associated with *Synechococcus* (Guedes et al., 2018) or more broadly with algae (Fullbright et al., 2018). Moreover, the *Gallionella* genus was previously observed colonizing filamentous algae (Mori et al., 2015).

Overall, these results show that the co-occurring surrounding community is, on average, more similar for more closely related *Synechococcus*. Members of this surrounding community could interact directly with *Synechococcus*, or share similarity in their abiotic niches.

## Discussion

In this study, we investigated *Synechococcus* niche preferences at relatively fine (sub-genus) taxonomic resolution using a 16S sequencing approach previously applied to other bacterial lineages (Koeppel and Wu, 2014; Tromas et al., 2018). However, we acknowledge that even at the finest possible taxonomic resolution, the 16S gene may be too slow-evolving to be a good marker for fast-evolving traits, such as toxin production (Berry et al., 2017) or bacteriophage resistance (Martiny et al., 2015). We found that water temperature, reservoir depth, and Nitrite concentration greatly affected the relative abundance of *Synechococcus* strains (Figure S6, S7). However, we did not find significant linear or non-linear relationships between genetic distance and preference for any of these abiotic niches. This was true when considering the whole range of genetic distance (>93% identity) or a restricted range of more closely related strains (>97%), suggesting that these traits are phylogenetically unconserved (at least within our dataset and at the resolution of the 16S marker). Such a lack of conservation would be expected for a trait such as Nitrite or Nitrate utilization in an organism capable of fixing nitrogen. Although some marine *Synechococcus* can fix nitrogen (Zehr et al., 2001), further investigation will be needed to determine if this is also the case in our freshwater community.

In contrast, total phosphorus (TP) preference was more phylogenetically constrained, even though it explained less variation in community structure than Nitrite (Figure S6, S7). Phosphorus use may therefore evolve “clock-like” along the *Synechococcus* phylogeny, following an approximately linear model in the range of 96-100% nucleotide identity (Figure 1). This contrasts with the observation in another cyanobacterium (*Prochlorococcus*) that organic phosphate niches are fast-evolving and thus correlated with the phylogeny only near the tips of the phylogenetic tree (Coleman & Chisholm 2010; Martiny et al., 2015). Consistent with this idea, we identified significant niche separation among very closely related (∼99% identical in 16S) *Synechococcus*, although the effect was driven by only three pairs of strains (Figure S12). We also note that TP is a fairly crude measurement, encompassing both organic and inorganic forms of phosphorus, and adaptation to these finer-grained niche dimensions may evolve differently in *Synechoccus.* TP may not even be the driving selective pressure and may simply be a marker of another important factor (*e.g.* runoff, overall biomass, etc.). Furnas Reservoir is meso-eutrophic and phosphorus is probably not limiting (concentration varies from 8 to 53µg/L, with a mean ∼25µg/L). This might explain why only 77 of the 780 *Synechococcus* strain pairs showed a significant co-response to phosphorus (Figure 1), indicating that the effect is driven by a small number of closely-related strains with similar co-responses, and cannot be generalized to all *Synechococcus*. This result might also be explained if *Synechococcus* represents a polyphyletic group (Coutinho et al., 2016), composed of different clades with different phosphorus niches. However, in our sample, *Synechococcus* is monophyletic (Figure S3, S4), excluding this explanation. Overall, our results demonstrate how abiotic factors can shape community composition within a genus but measured abiotic niches generally do not evolve along the genus-level phylogeny.

Habitat filtering was originally defined to describe how abiotic factors select for genetically similar (more closely related) organisms (Cadotte and Tucker 2017). Yet in practice, it is difficult to disentangle the contributions of biotic and abiotic factors, and thus to satisfactorily define habitat filtering (Kraft et al., 2015). In this study, we therefore estimated the influence of both unmeasured abiotic variables and biotic factors, and found a negative association with genetic distance (Figure 2). This suggests that more closely related *Synechococcus* tend to share similar realized niches. As a result, we confirmed that genetically similar *Synechococcus* nodes tend to co-occur, (Figure 3), as observed in studies using genomic (rather than 16S) similarity (Kamneva, 2017). This result could be unintuitive especially if *Synechococcus* nodes co-occur, but do not share abiotic dimensions. For example, two taxa could be present in the same samples, but at very different relative abundances. Therefore, they would co-occur but respond quantitatively to different environmental variables. Overall, this result is consistent with habitat filtering, broadly defined to include both biotic and abiotic factors.

Here we found that commonly-measured abiotic environmental factors (*e.g.* nutrients, temperature) may be insufficient to capture relevant niche dimensions, and biotic co-occurrence networks could be more informative about the evolution of niche preferences. In this study, we measured abiotic factors such as temperature and nutrients, which are important in structuring microbial communities, and yet these factors are largely insufficient to account for realized niches or their distribution across the *Synechococcus* phylogeny. However, more closely-related *Synechococcus* tend to co-occur with more similar surrounding microbial communities. As previously suggested, the surrounding microbial community could provide hints as to the nature of a realized niche (Cohan and Koeppel, 2008; Tromas et al., 2018). For example, the presence of certain taxa could indicate the importance of an unmeasured abiotic factor (*e.g* Acidophiles or Alkaliphiles as indicators of pH). In some cases, co-occurrence may indicate real biological interactions (*e.g.* physical association, predation, cross-feeding, etc.). Some of the bacteria whose co-occurrence with *Synechococcus* was strongly phylogenetically conserved have previously been found in physical association with algae and cyanobacteria (Table S1; Mori et al. 2015; Fullbright et al., 2018) and future work could experimentally test the nature of possible interactions with *Synechococcus* strains. Alternatively, co-occurring microbes could simply share similar unmeasured niches (including unmeasured abiotic or biotic factors such as phage or protozoan grazers), and mining their genome sequences or metabolic capabilities could provide insight into the nature of these niches. In this case, considering biotic factors allowed us to uncover a signal of habitat filtering within *Synechococcus* that would have been obscured by only considering commonly measured abiotic factors.

## Supporting information

Environmental variables Table

## Data availability

Raw sequence data have been deposited NCBI GenBank under BioProject number PRJNA544938.

## Acknowledgments

NT is funded by a project from the European Union’s Horizon 2020 research and innovation program under the Marie Sklodowska-Curie grant agreement No 656647. BJS was supported by a Canada Research Chair and NSERC Discovery Grant. Sampling and samples processing were supported by a grant from Furnas Centrais Hidroelétrica S.A. (Brazil) to AG. JSMP and DAP received a scholarship from CAPES (Coordenação de Aperfeiçoamento de Pessoal de Nível Superior). A sabbatical visit of AG to the Université de Montréal AG was supported by a CAPES Senior Fellowship.

