## Supplementary Files for "The evolution of realized niches within freshwater *Synechococcus*"

**SUPPLEMENTARY FIGURES**

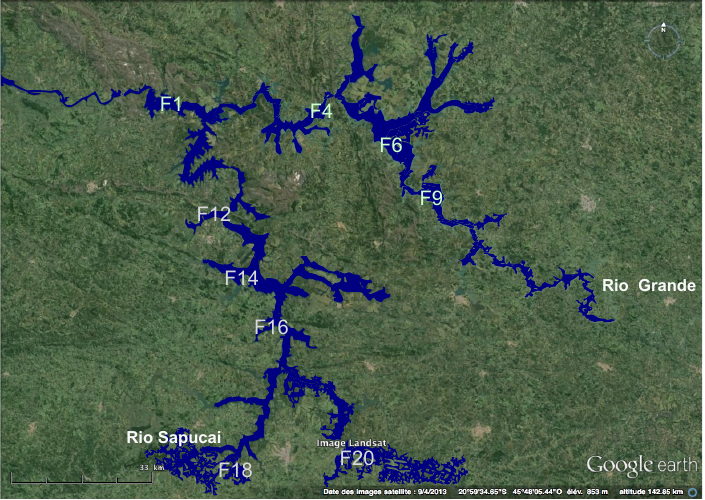

**Figure S1. Map of the sampling stations. (Google Earth).** Furnas reservoir is located in South-Eastern Brazil (20o40’S; 46o19’W). This large reservoir (1,440 Km^2^; 20.95 billion m^3^) is composed of two major rivers, Grande and Sapucaí, and several small tributaries. Stations F12-F20 are mainly located in regions occupied by livestock, agricultural activities and several small to medium cities. Stations F4 –F9 are located in less impacted areas. Station F1 is located close to the dam, at the intersection of both rivers.

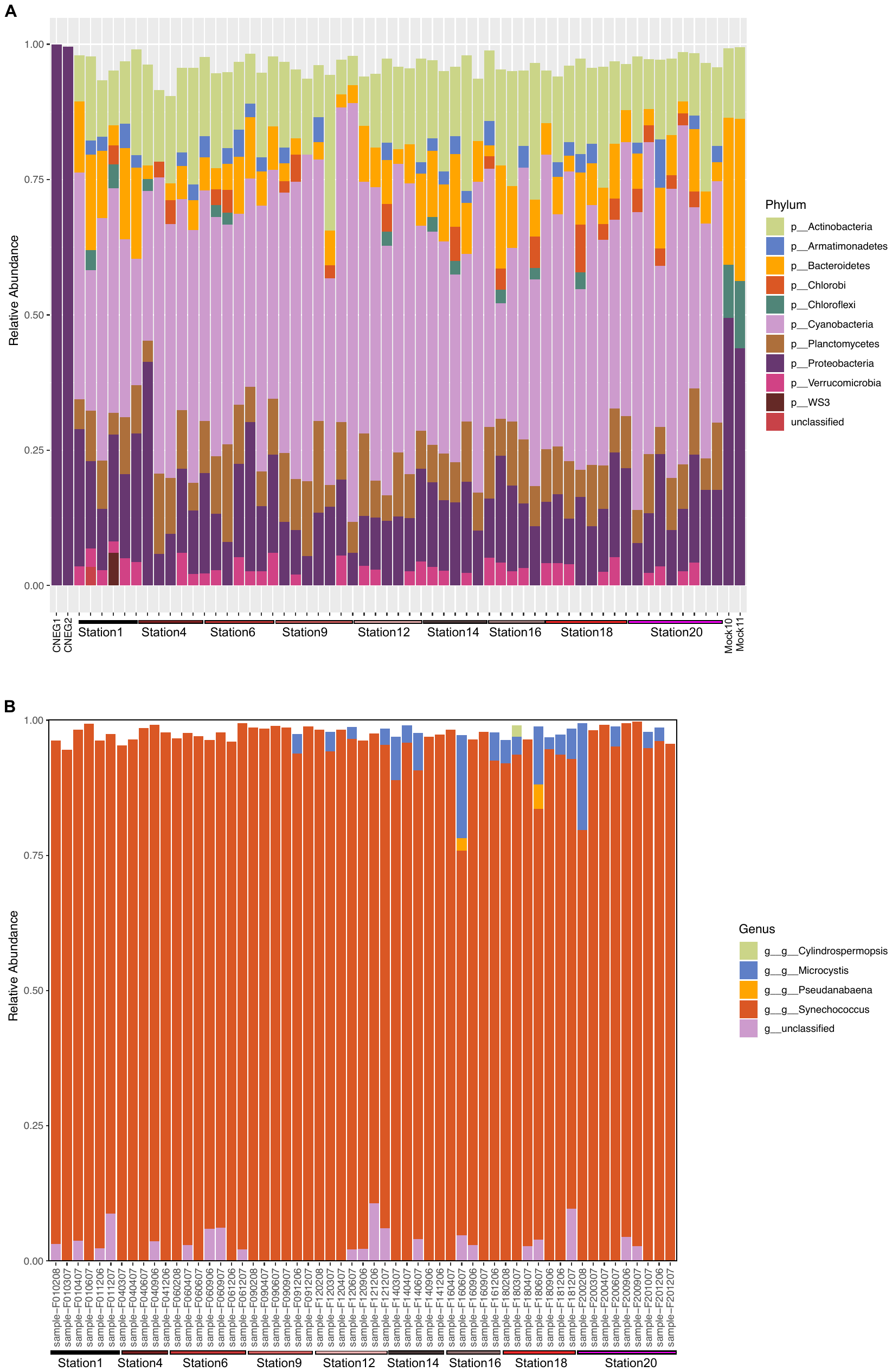

**Figure S2. Barplot of the relative abundance of microbial taxa across stations F4-F9 and F12-20 of Furnas reservoir. A.** Barplot of the microbial composition at the phylum level, including negative controls (CNEG1 and CNEG2; left) and mock communities (Mock10 and Mock 11; right). **B.** Barplot of the cyanobacterial composition at the genus level.

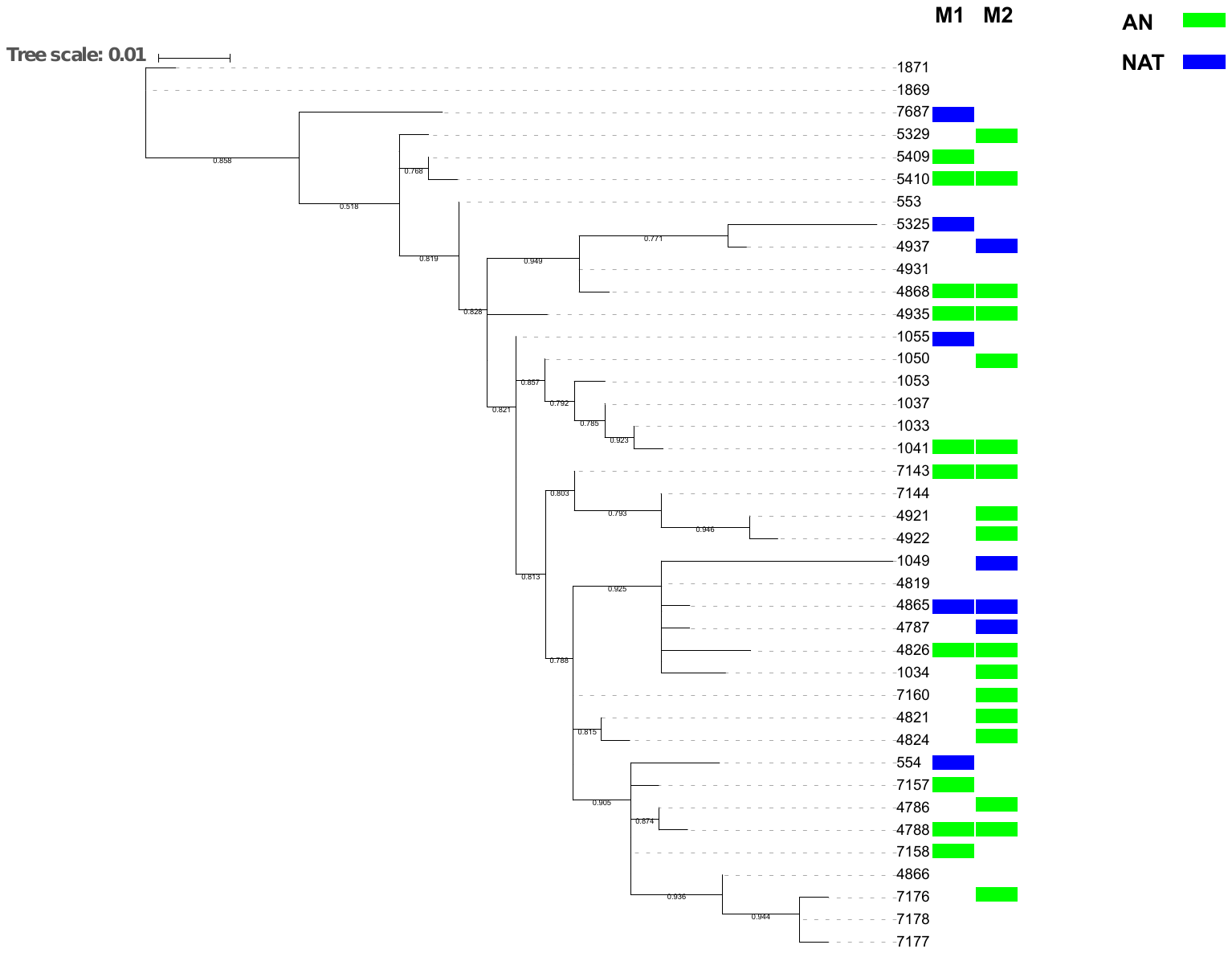

**Figure S3. Phylogenetic distribution of *Synechococcus* nodes across the less impacted (NAT, F1-F9) and more impacted (AN, F12-20) branches of the reservoir.** Two different methods (ALDEx2; M1 and DESeq2; M2) were used to determine if a *Synechococcus* node is differentially associated with one of these two branches.

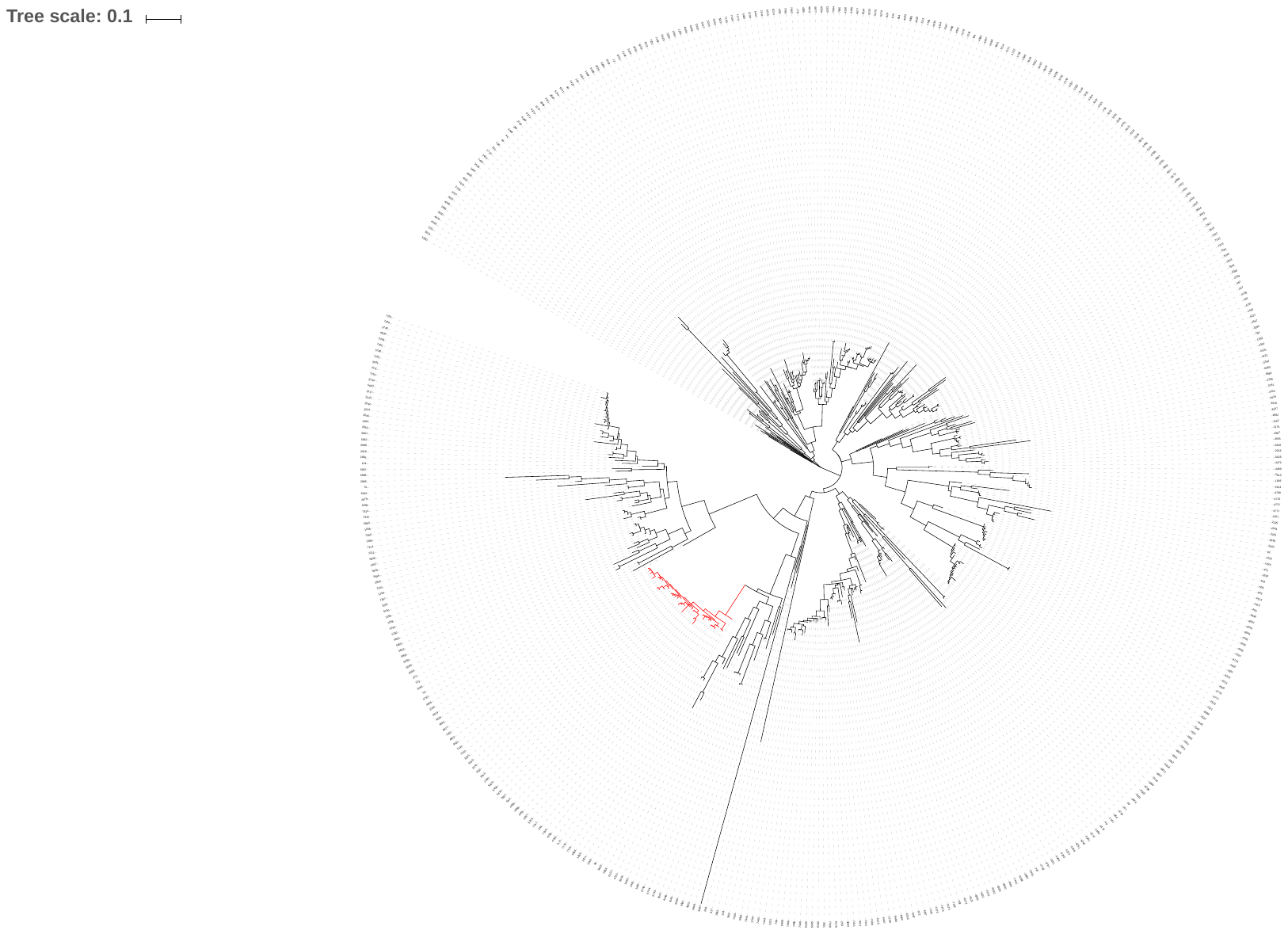

**Figure S4. Phylogenetic characterization of the 16S rRNA gene from the Furnas reservoir.** *Synechococcus* nodes are highlighted in red, forming a monophyletic group.

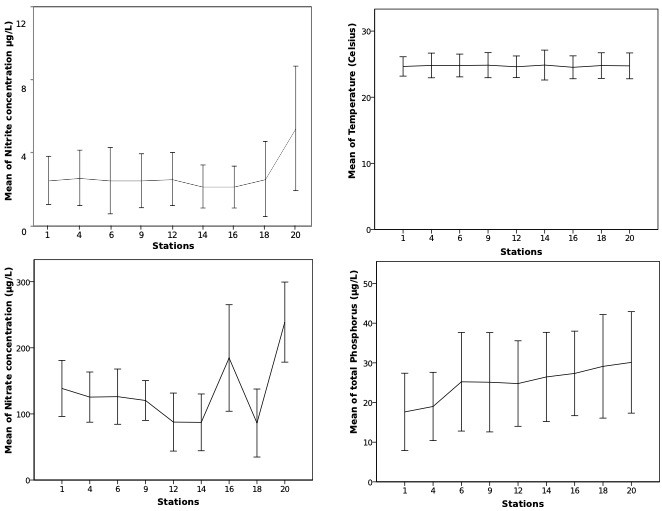

**Figure S5. Geographical variation (across stations) of different abiotic variables measured in Furnas reservoir from 2006-2009.** Error bars correspond to the standard deviation and represent temporal variability (years and months) both across the season and among years sampled.

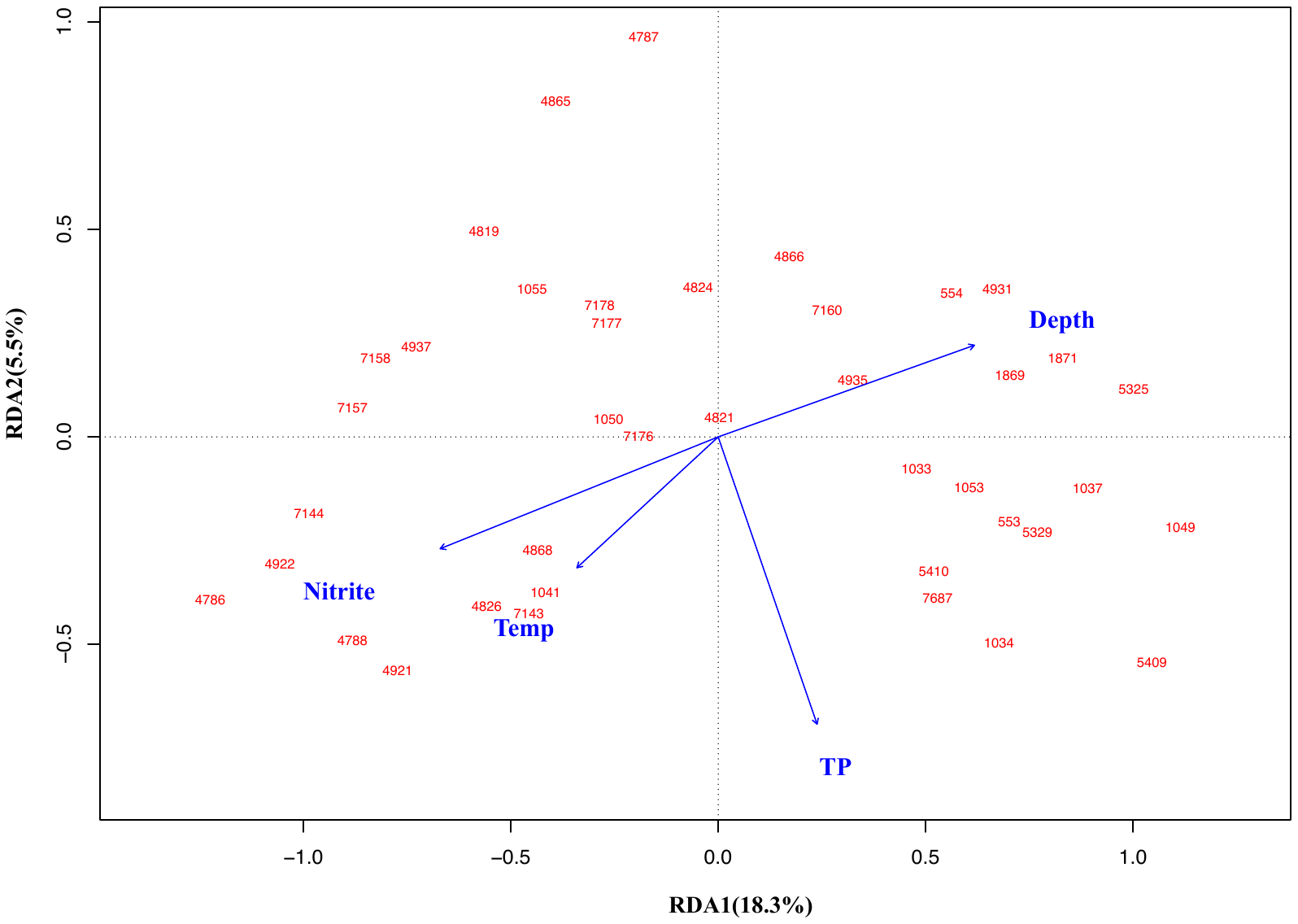

**Figure S6. Parsimonious redundancy analysis of the *Synechococcus* community versus** **the forward-selected environmental variables.** Forward selection identified Nitrite, total phosphosrus (TP), depth (ZE), and water temperature (Temp) as important variables. The first two constrained axes (RDA 1 and RDA 2) explained 23.8% of the variance observed in the *Synechococcus* community composition. *Synechococcus* MED node IDs are represented in red.

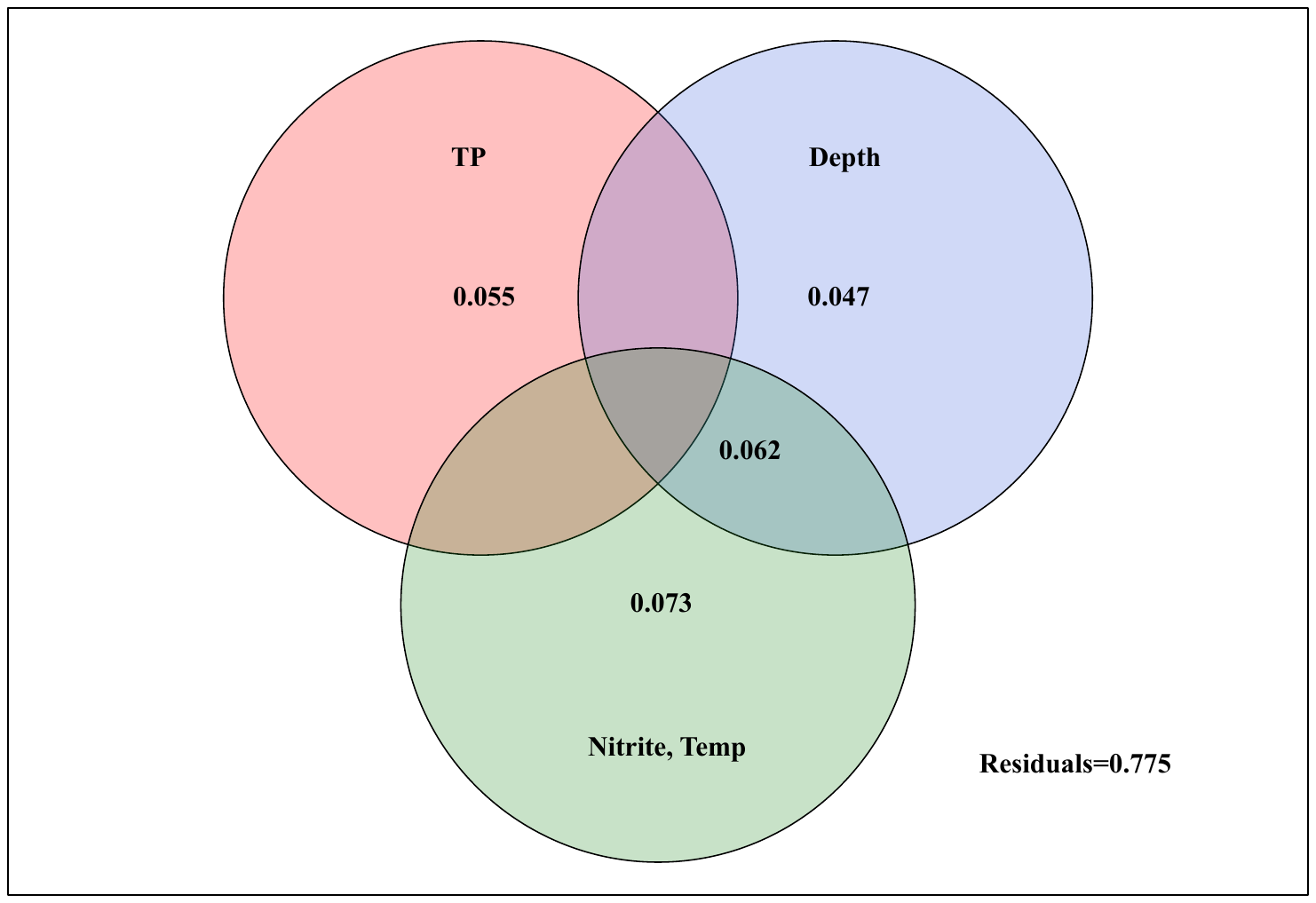

**Figure S7. Variance partitioning analysis of the forward-selected environmental variables.** Based on the RDA (Fig. S6) we lumped groups of explanatory variables that best separated the first two RDA dimensions (i.e., group 1 = TP; group 2 = Depth; group 3 = Temp, Nitrite) and tested the proportion of variation in *Synechococcus* community composition that was jointly and uniquely explained by each group of variables. Temperature and Nitrite alone explained the greatest, unique amount (7.3%) of the *Synechococcus* community.

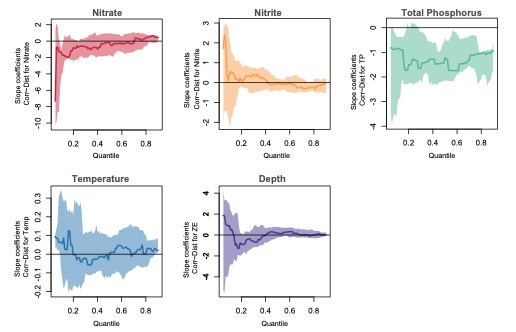

**Figure S8. Significance plot of the quantile regression models of co-response vs genetic distance.** Quantile slope estimates and corresponding confidence intervals (bands) of the LVM co-responses vs genetic distance across all quantiles of genetic distance. Note that slopes at the 5^th^, 10^th^, 15^th^, 25^th^, 35^th^, 45^th^, 55^th^, 65^th^, 75^th^, 85^th^ and 90^th^ quantiles are shown in Figure 2. Significant slopes are those where both the slope estimate and confidence band do not overlap with zero.

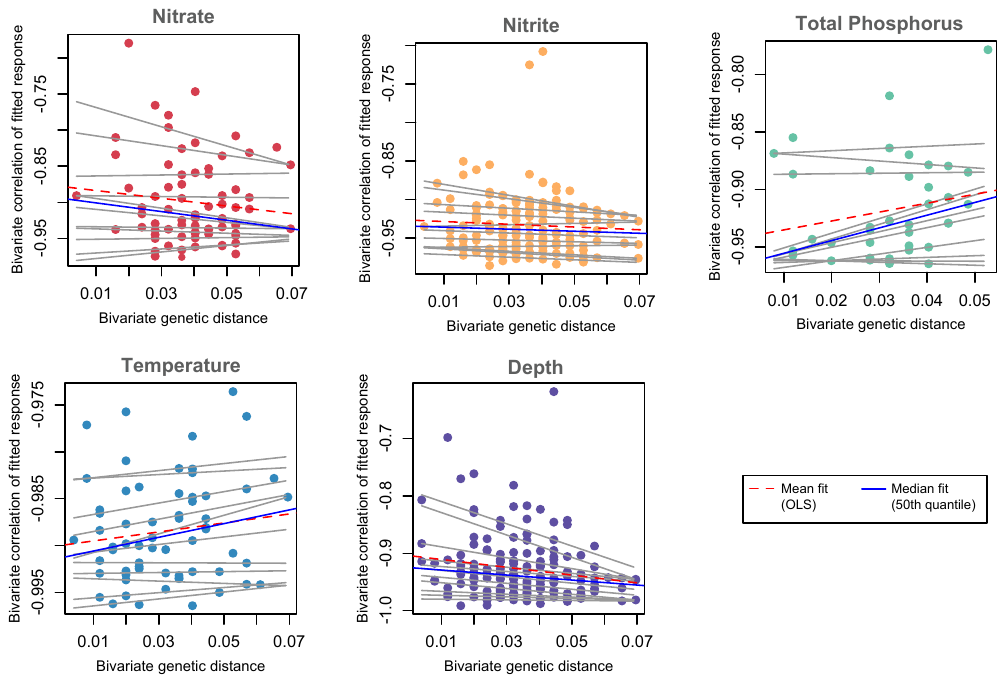

**Figure S9. Scatterplot and quantile regression models of niche separation (negative correlation among LVM fitted values) vs. genetic distance.** See Figure 1 for details.

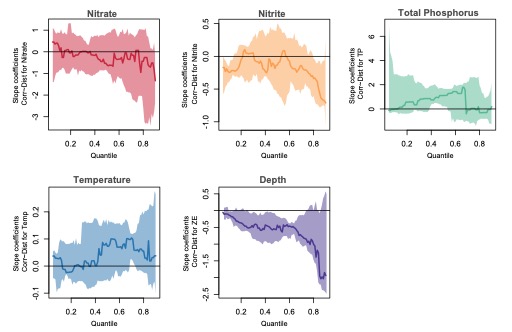

**Figure S10.** **Significance plot of the quantile regression models of niche separation vs genetic distance.** See Figure S8 for details.

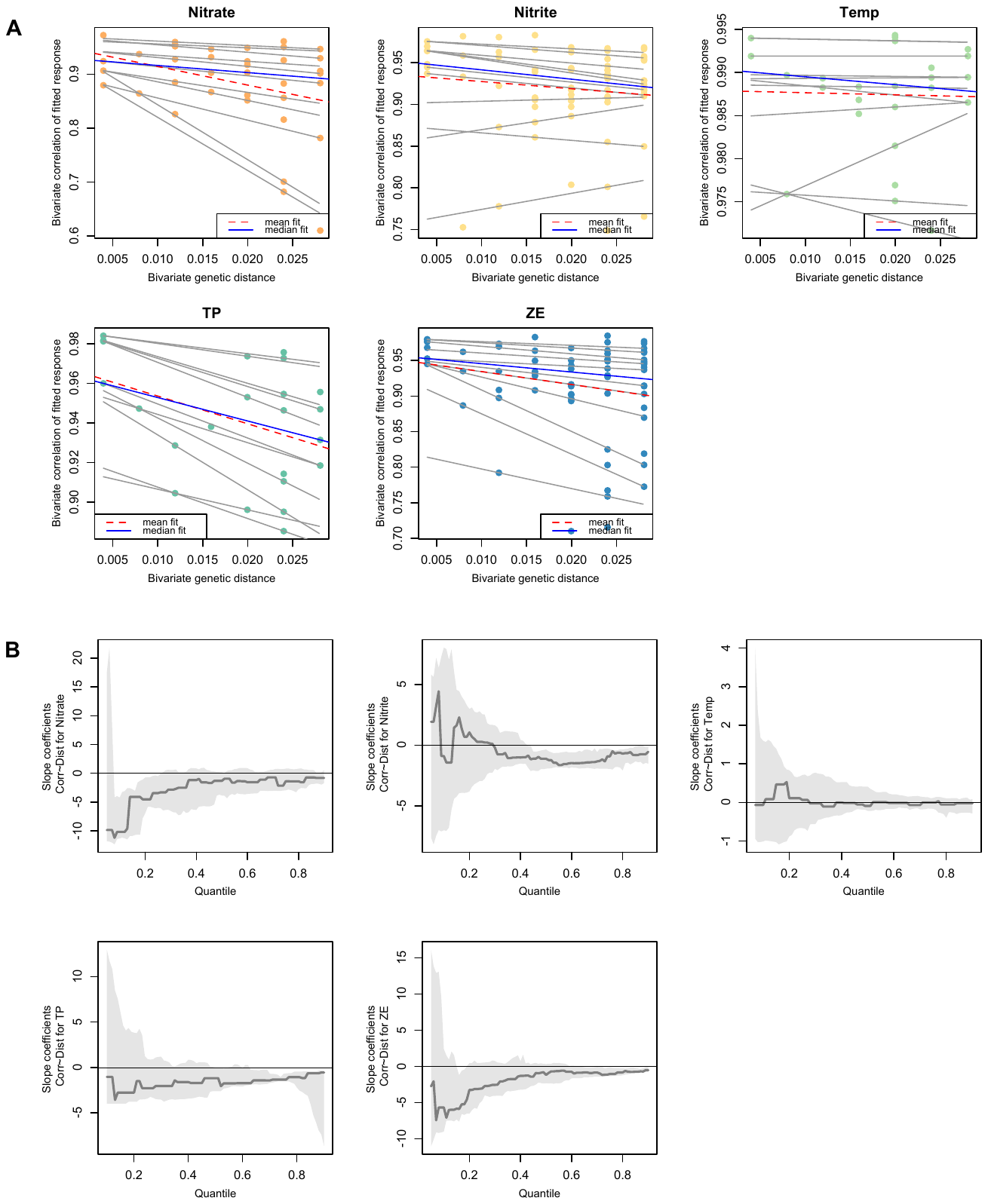

**Figure S11. Scatterplot and quantile regression models of niche similarity.** (A) A similar analysis as Figure 1 was performed in the restricted range of 100-97% identity. The bottom panels (B) show significance plots, equivalent to those shown in Figure S8, but in a restricted range of genetic identity.

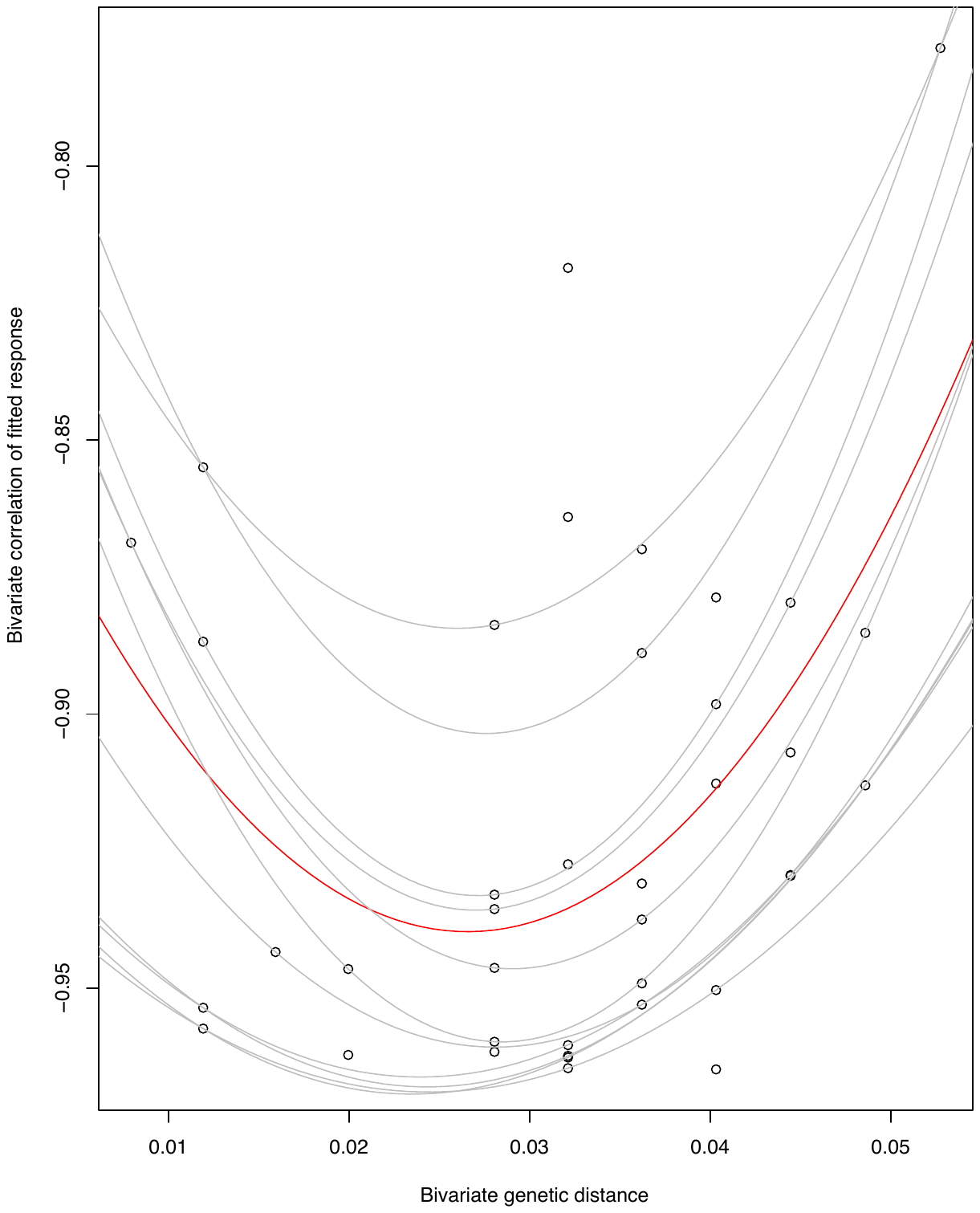

**Figure S12. Scatterplot and quantile regression models of niche separation for TP (negative correlation among LVM fitted values) vs. genetic distance.** Shown are the scatterplots of LVM negative co-responses versus each variables superimposed with the lines for 5^th^, 10^th^, 15^th^, 25^th^, 35^th^, 45^th^, 55^th^, 65^th^, 75^th^, 85^th^ and 90^th^ quantile regression fits and the mean fit (red line).

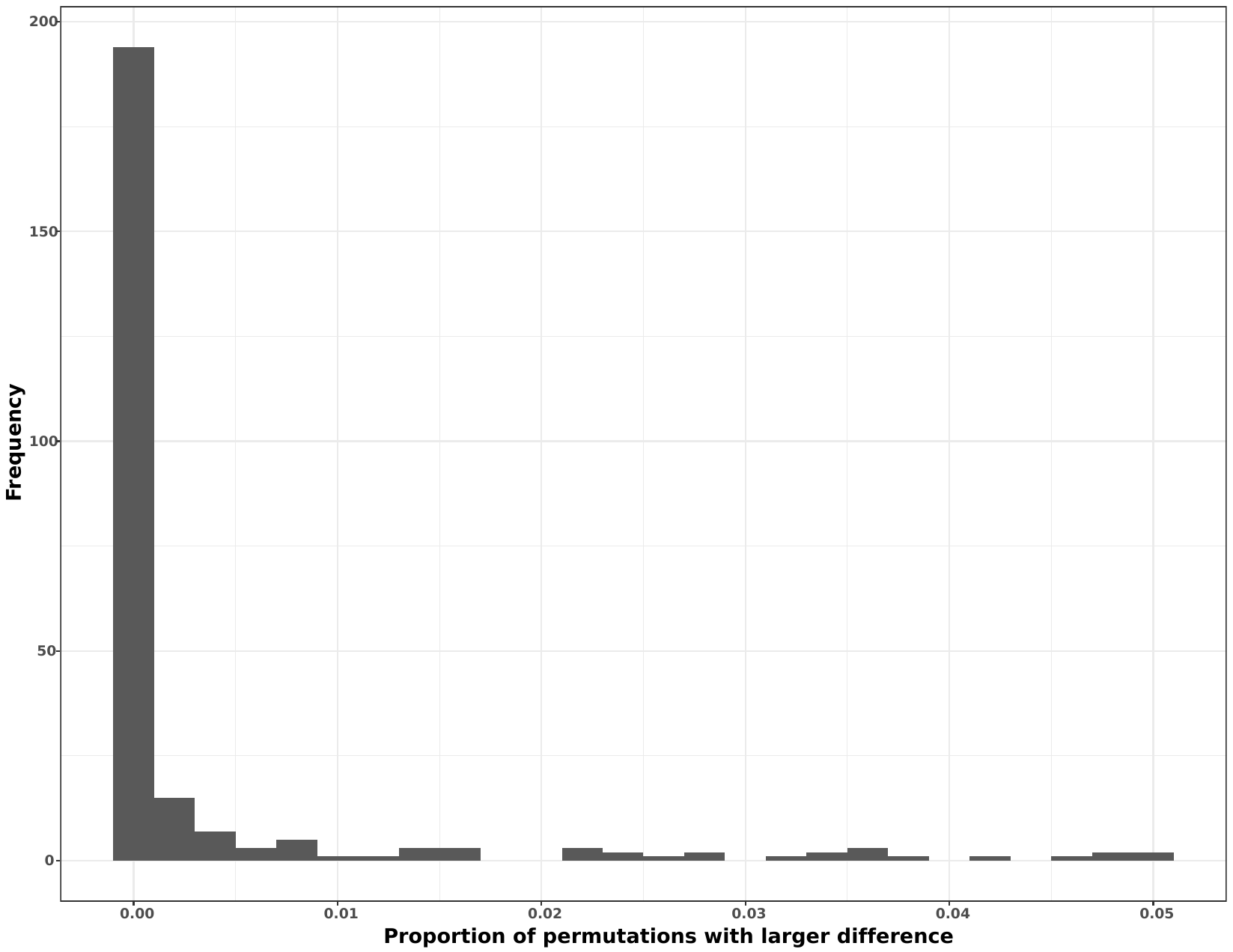

**Figure S13. Permutation test of the association between *Synechoccus* genetic distance and association with similar surrounding communities.** We estimated the proportion (*p*) of false positive correlations between genetic distance of *Synechococcus* nodes and the co-occurrence with non- *Synechococcus* taxa. The distribution is based on 1000 permutations for each of 253 non-*Synechococcus* taxa, randomizing the association between |Δ*r*| and genetic distance (Methods and R_script2). We then calculated *p* as the number of permutations yielding a larger correlation than observed (numerator) divided by the total number of permutations (denominator).
